# Biophysical modeling of the electric field magnitude and distribution induced by electrical stimulation with intracerebral electrodes

**DOI:** 10.1101/2023.01.13.523921

**Authors:** Fabiola Alonso, Borja Mercadal, Ricardo Salvador, Giulio Ruffini, Fabrice Bartolomei, Fabrice Wendling, Julien Modolo

## Abstract

Intracranial electrodes are used clinically for diagnostic or therapeutic purposes, notably in drug-refractory epilepsy (DRE) among others. Visualization and quantification of the energy delivered through such electrodes is key to understanding how the resulting electric fields modulate neuronal excitability, i.e. the ratio between excitation and inhibition. Quantifying the electric field induced by electrical stimulation in a patient-specific manner is challenging, because these electric fields depend on a number of factors: electrode trajectory with respect to folded brain anatomy, biophysical (electrical conductivity / permittivity) properties of brain tissue and stimulation parameters such as electrode contacts position and intensity.

Here, we aimed to evaluate various biophysical models for characterizing the electric fields induced by electrical stimulation in DRE patients undergoing stereoelectroencephalography (SEEG) recordings in the context of pre-surgical evaluation. This stimulation was performed with multiple-contact intracranial electrodes used in routine clinical practice. We introduced realistic 3D models of electrode geometry and trajectory in the neocortex. For the electrodes, we compared point (0D) and line (1D) sources approximations. For brain tissue, we considered three configurations of increasing complexity: a 6-layer spherical model, a toy model with a sulcus representation, replicating results from previous approaches; and went beyond the state-of-the-art by using a realistic head model geometry.

Electrode geometry influenced the electric field distribution at close distances (~3 mm) from the electrode axis. For larger distances, the volume conductor geometry and electrical conductivity dominated electric field distribution. These results are the first step towards accurate and computationally tractable patient-specific models of electric fields induced by neuromodulation and neurostimulation procedures.

## Introduction

Stereoelectroencephalography (SEEG) is routinely used to identify epileptogenic zones (EZ) in patients with drug-refractory epilepsy (DRE) who are potential candidates to surgery. This technique consists in the surgical implantation of intracranial depth multiple-lead electrodes that include 10-15 contacts. Both recordings of spontaneous activity and electrical stimulation *via* SEEG electrodes are used to determine the 3-dimensional spatiotemporal organization of the brain epileptogenic network, and identify potential targets for surgical resection [1].

During pre-surgical evaluation, electrical stimulation is routinely used to probe different brain regions spatially sampled by electrode contacts to obtain a functional map where epileptogenic dysfunctional cortical regions are distinguished from functional ones based on their electrophysiological responses. Typically, the presence of a stimulation-induced post-discharge that can possibly trigger clinical symptoms and seizures is indicative of an epileptogenic region [2]. In addition, local, bipolar stimulation has been reported as a method to probe recorded brain regions and estimate their level of excitability using a quantitative index referred to as the Neural Network Excitability Index [3]. Furthermore, therapeutic brain stimulation using SEEG-like electrodes, similar to those used in deep brain stimulation (DBS) in Parkinson’s disease, has been investigated as a possibility to decrease pathological hyperexcitability of epileptogenic regions [4]. However, one major challenge is that the characteristics of induced electric fields (magnitude, orientation) in brain tissues, and its anatomical targets are not always clearly characterized. Such uncertainties limit the mechanistic understanding of stimulation effects on brain tissues (e.g., grey/white matter). Furthermore, dosimetric evaluations of the *in situ* electric field are typically computationally extensive, motivating the development of alternative methods that would provide faster while still reliable dosimetric estimates. Consequently, quantifying the electric field induced by SEEG electrodes during electrical stimulation is of considerable interest to further develop and optimize diagnostic and therapeutic applications in DRE in particular, as well as in other neurological disorders using similar DBS intracranial electrodes like dystonia, essential tremor, obsessive-compulsive disorders and Parkinson’s disease, to name a few.

From the biophysics viewpoint, in biological tissue, the electric field distribution is governed by a partial differential equation (as per Maxwell equations of bioelectromagnetism) that can be solved analytically (in principle at least, since this can prove extremely challenging if possible at all depending on the system investigated), approximating the electrodes as point sources; or numerically using the finite element method (FEM) for realistic models of the electrode and surrounding brain tissue. Regarding brain tissue, the use of realistic head models based on magnetic resonance imaging (MRI) data including several tissue types significantly increases the complexity of integrating geometrically-accurate representations of electrodes, resulting in models that are computationally expensive and technically challenging to implement. More specifically, a major problem with image-based modeling relates to the generation of a proper mesh for the connection between two complex geometries, such as in the case of intracranial electrodes inserted in brain tissue. Therefore, possible alternatives to overcome this problem consist in approximating the 3D electrode’s cylindrical shape by a point (0D) or a line (1D) source. Therefore, using realistic and simplified head models, the objective of this study was to evaluate if 0D and 1D current source approximations are sufficiently accurate to quantitatively characterize the electric field induced by electrical stimulation applied with SEEG electrodes, in comparison with a more realistic 3D model of SEEG electrodes. By replicating previous results from the literature, and also providing novel results in the context of realistic head models, our study provides guidelines for the biomedical engineering community regarding modeling compromises regarding electric fields induced by standard, intracranial electrodes.

## Results

Results obtained for the 0D and 1D approximation are reported in Fig. 1 for the spherical model (Fig. 1a). As depicted in Fig. 1b-d, the electric field (abbreviated as “E-Field” hereafter) estimation was qualitatively similar for the three electrode models (realistic cylindrical geometry source, line source, point source). Nevertheless, some differences were observed and are displayed in Fig. 1eh. When comparing specific electric field isovalues at a plane crossing the electrode (Fig. 1e), the line source approximation better matched the E-Field shape obtained with the cylindrical model.

**Figure 1.**
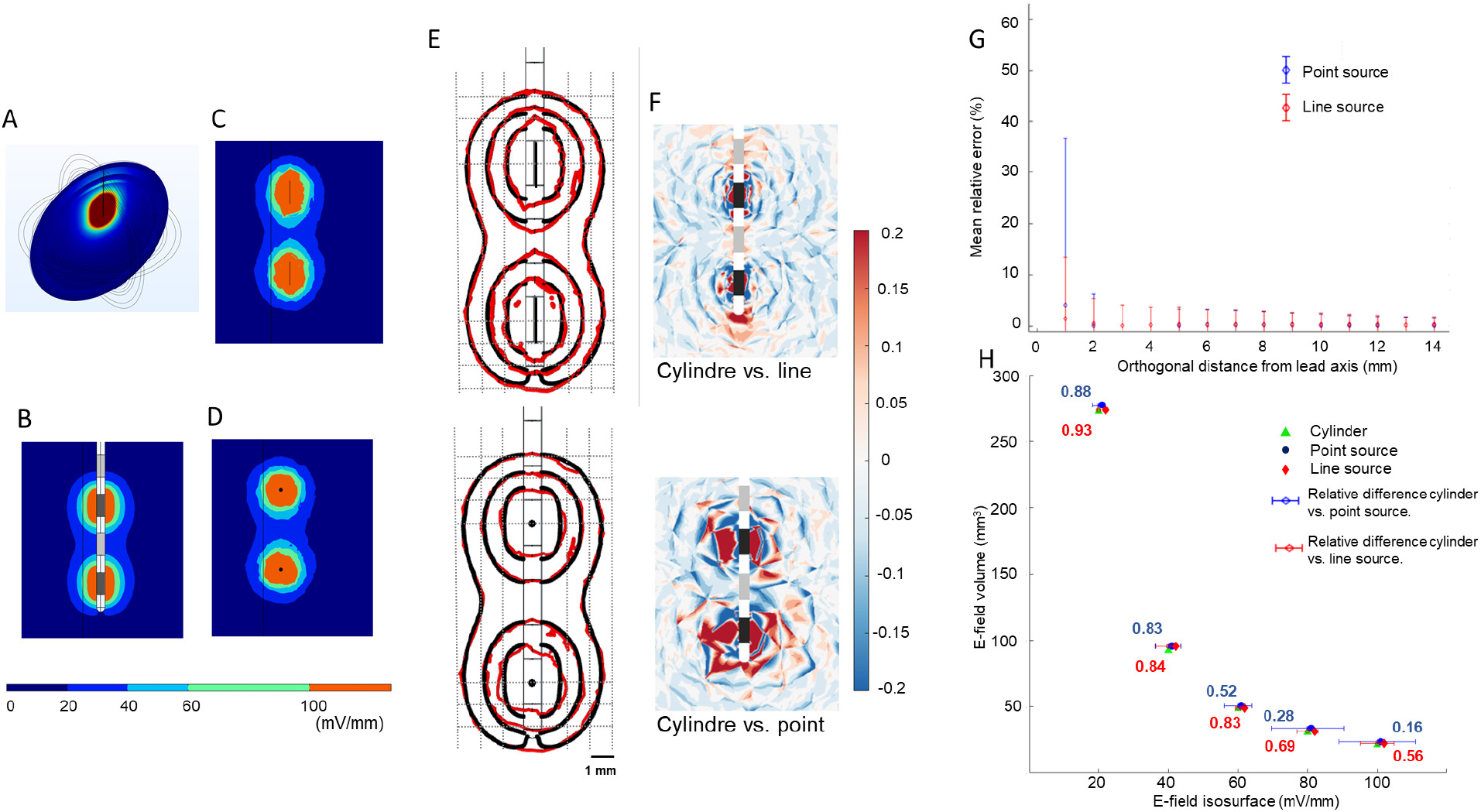
0D and 1D E-Field approximation, spherical volume conductor model. A) Electric field distribution at the plane crossing the electrode for a non-adjacent configuration injecting ±1 mA with the contact pair. B) Electric field for the realistic model, C) line source approximation, and D) point source approximation. E) Isofield lines (20, 40, 100 mV/mm) for the realistic model (black lines), line source approximation (top, red lines), and point source approximation (bottom, red lines). F) Relative error, calculated on the same plane, between the realistic and line approximation (top) and point source approximation (bottom). G) Mean relative error and standard deviation between the approximations and realistic model of the electric field as a function of the orthogonal distance from the electrode axis. H) Electric field volume within the 20, 40, 60, 80, 100 mV/mm isosurfaces for the three electrode models along with the corresponding relative error bars and Dice coefficient for the point (in blue) and the line (red) source.

The isocontours for the point source were more “rounded”, while those for the line source approximated better the elongated shape of the isocontour obtained with the cylindrical electrode. The relative error displayed in Fig. 1f showed an overestimation of the E-Field at the level of active contacts that was more pronounced for the point source as compared to the line source. As shown in Fig. 1g, the error was lower for the line source approximation, regardless of the distance with respect to the electrode. Let us mention that both approximations slightly overestimated the electric field magnitude for the volume comparison. The relative difference between line source and cylindrical source (error bars in Fig. 1h) ranged from 0.4% to 5% for the 20 and 100 mV/mm isosurfaces, respectively; while for the point source the relative difference ranged between 2% and 11%. The similarity between the volumes, as quantified by the Dice coefficient (eq. 1), showed a better performance of both approximations at lower E-Field magnitudes. For large E-Field magnitudes, however, the line source resulted in a higher Dice coefficient (i.e., performed better).

Results obtained for the 0D and 1D approximations are reported in Fig. 2 in the case of the toy model, in which a sulcus is represented and when the electrode is inserted parallel to the sulcus at a distance of 5.1 mm. Fig. 2a presents the E-Field distribution at the plane crossing the electrode.

**Figure 2.**
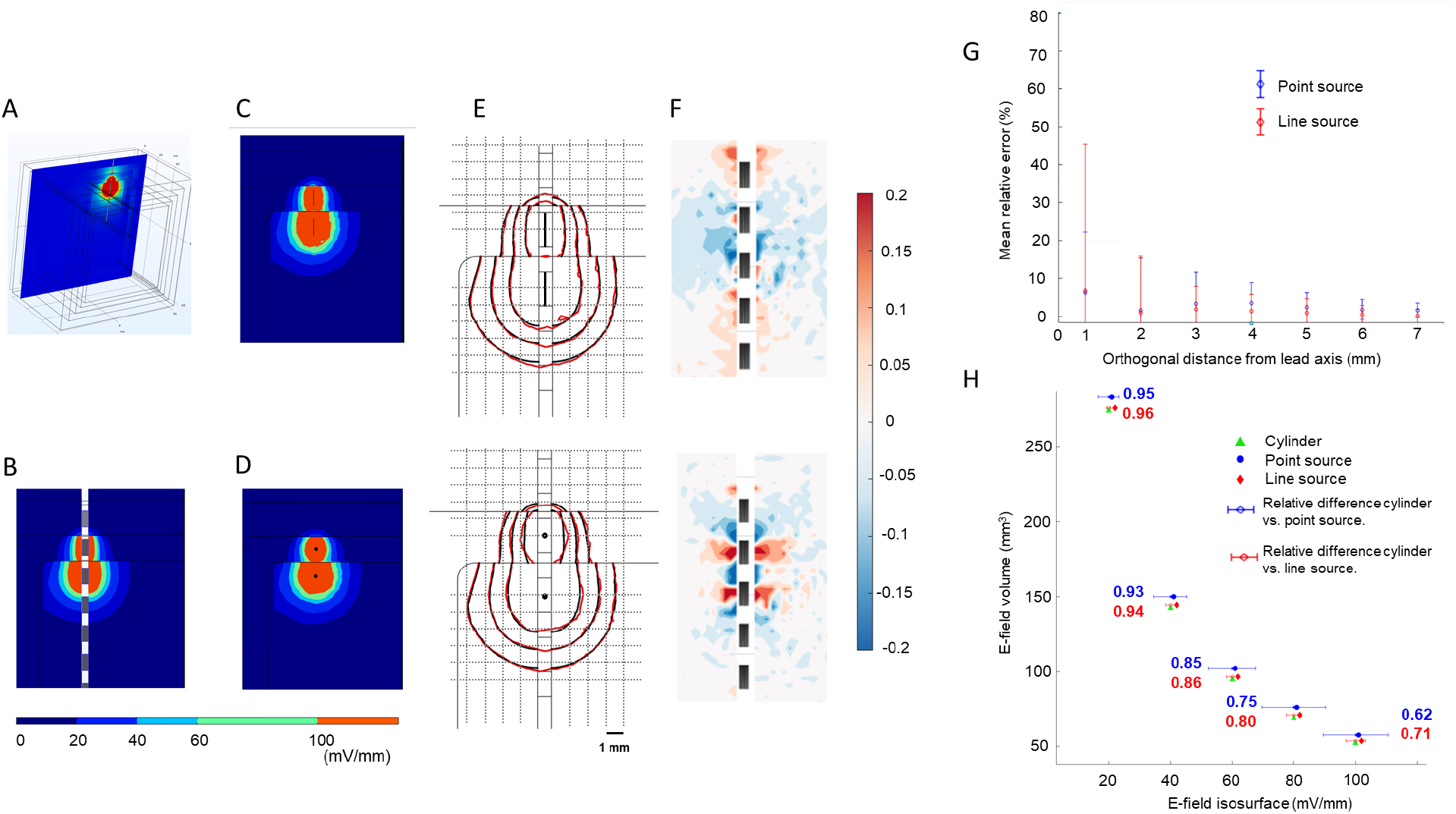
0D and 1D E-Field approximation, toy volume conductor model, with the electrode parallel to the sulcus. A) Electric field distribution at the plane crossing the electrode for a non-adjacent configuration injecting ± 1 mA with the contact pair. B) Electric field for the realistic model, C) line source approximation, and D) point source approximation. E) Isofield lines (20, 40, 100 mV/mm) for the realistic model (black lines), line source approximation (top, red lines) and point source approximation (bottom). F) Relative error, calculated on the same plane, between the realistic and line approximation (top) and point source approximation (bottom). G) Mean relative error and standard deviation between the approximations and realistic model of the electric field as a function of the orthogonal distance from the electrode axis. H) Electric field volume within the 20, 40, 60, 80, 100 mV/mm isosurfaces for the three electrode models showing the corresponding relative error bars and the Dice coefficient (blue for the point, and red for the line source).

For the toy model, when placing the electrode parallel to the sulcus, results were qualitatively similar to those obtained with the spherical model except that discontinuities were observed in the E-Field spatial distribution (Fig. 2b-d), and were not present in the spherical model. As discussed below, this phenomenon can be explained by abrupt changes in conductivity values corresponding to white and gray matter. With one of the active contacts in the white matter and the other in the gray matter, the electric field is distributed farther away due to the lower electrical conductivity of white matter.

The isocontours displayed in Fig. 2e illustrate a better performance from the line source approximation by presenting a more elongated shape that better matches the field obtained with the cylindrical source. The point source, in contrast, resulted in a more rounded shape which was also revealed by the relative error showing an overestimation of E-field magnitude at the middle of the contact (Fig. 2f, bottom panel). The line source had a slightly higher relative error than the point source (Fig. 2g) at a distance of 1 mm, but decreased beyond 2 mm from the electrode axis. The E-Field volume computed for both approximations was larger than that obtained by the cylindrical model, and this difference was larger for higher values of the electric field, i.e. 100 V/m which corresponds to the magnitude closer to the electrode (Fig. 2h). The relative difference between volumes ranged between 0.7% to 3% for the line source, and between 3% to 10% for the point source. The Dice coefficient (shown to the side of the error bars in Fig. 2h) showed a good performance (~0.95) for both approximations for E-Field magnitudes of 20 and 40 mV/mm. The Dice coefficient decreased for larger E-Field magnitudes, falling to 0.7 and 0.6 for the line and point source approximations, respectively.

When the electrode was tilted and therefore crossed layers with different conductivities, the line source approximated more accurately the E-field obtained by the realistic electrode representation. The qualitative comparison of the EF at the plane traversing the shaft (Fig. 3a) showed a very similar electric E-Field for both approximations (Fig. 3b-d). Using the non-adjacent configuration, both active electrodes were placed in gray matter, thus showing a similar extension of the E-field around each electrode. The isocontours, in Fig. 3e, showed a better performance of the line source approximation. The relative error at the plane presented in Fig. 3f showed an underestimation of both approximations between the active contacts, i.e., the region filled with CSF. The point source however, showed a lower relative error as a function of the orthogonal distance (Fig. 3g), indicating a better performance closer to the electrode (between 1 and 3 mm). In contrast to the previous cases, the EF volume difference (Fig. 3h) was similar between the point and line source approximations ranging both between ~3% to ~8%. This was also reflected by the Dice coefficient, except for the volume at 100 mV/mm where the line source performed slightly better (~0.7) as compared to the point source (~0.4).

**Figure 3.**
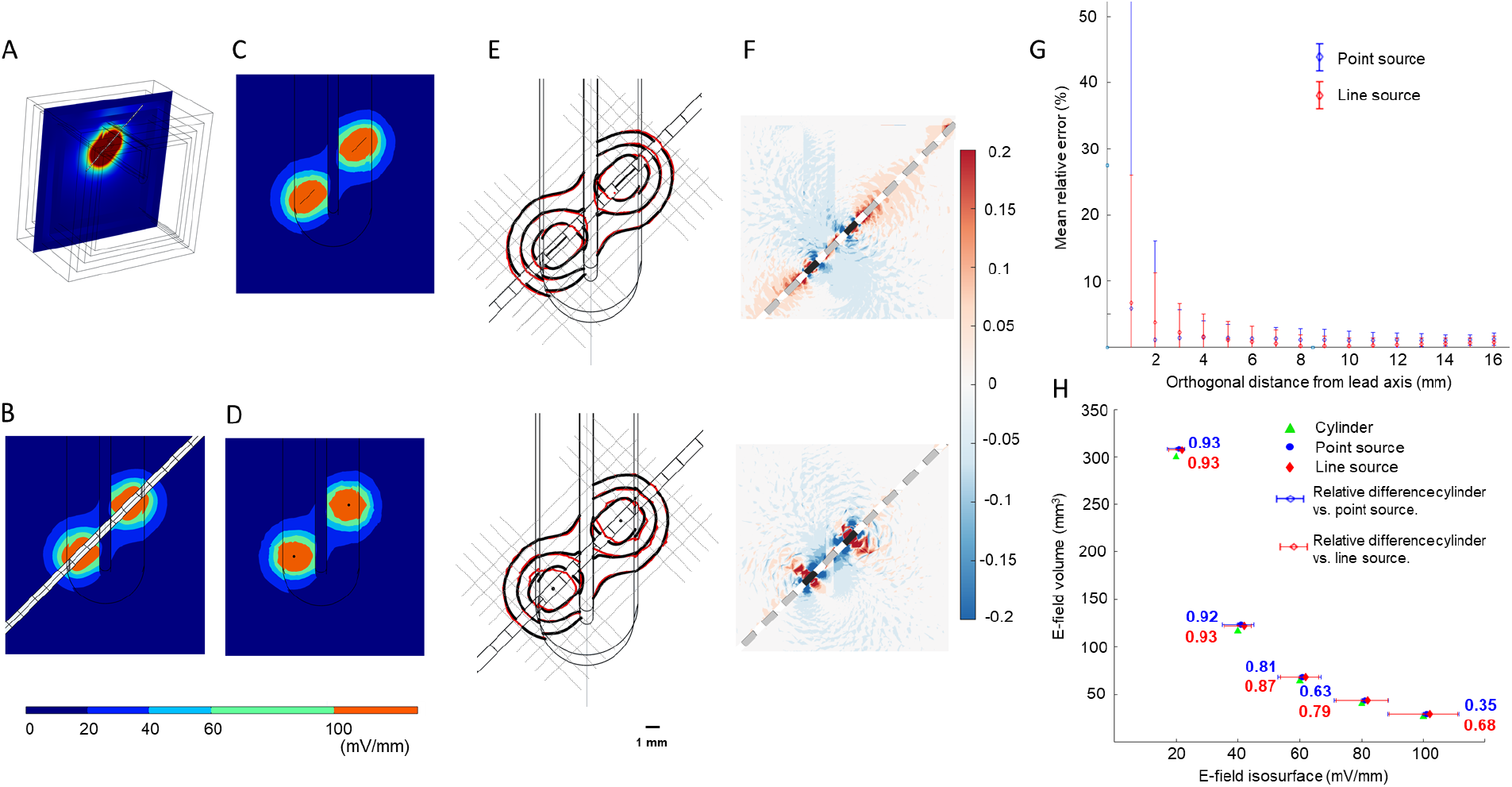
Toy model with a tilted electrode. A) Electric field distribution at the plane crossing the electrode for a non-adjacent configuration injecting ± 1 mA with the contact pair. B) Electric field for the realistic model, C) line source approximation, and D) point source approximation. E) Isofield lines (20, 40, 100 mV/mm) for the realistic model (black lines), line source approximation (top, red lines) and point source approximation (bottom). F. Relative error, calculated on the same plane, between the realistic and line approximation (top) and point source approximation (bottom). G) Mean relative error and standard deviation between the approximations and the realistic model of the electric field as a function of the orthogonal distance from the electrode axis. H) Electric field volume within the 20, 40, 60, 80, 100 mV/mm isosurfaces for the three electrode models along the corresponding relative error bars and the Dice coefficient shown in blue for the point and in red for the line source.

For the realistic head model, electric field distribution was clearly impacted by the electrical conductivity of surrounding medium. At the level of contact C3, the electric field magnitude of 20 mV/mm for instance, both electrode models (cylindrical and point source approximations) showed a very similar shape. (Fig. 4a-b). In contrast, for contact C1 (surrounded by CSF which has higher conductivity), the electrode geometry dominated the E-Field distribution, showing the elongated and circular shape of the cylinder and point sources, respectively.

**Figure 4.**
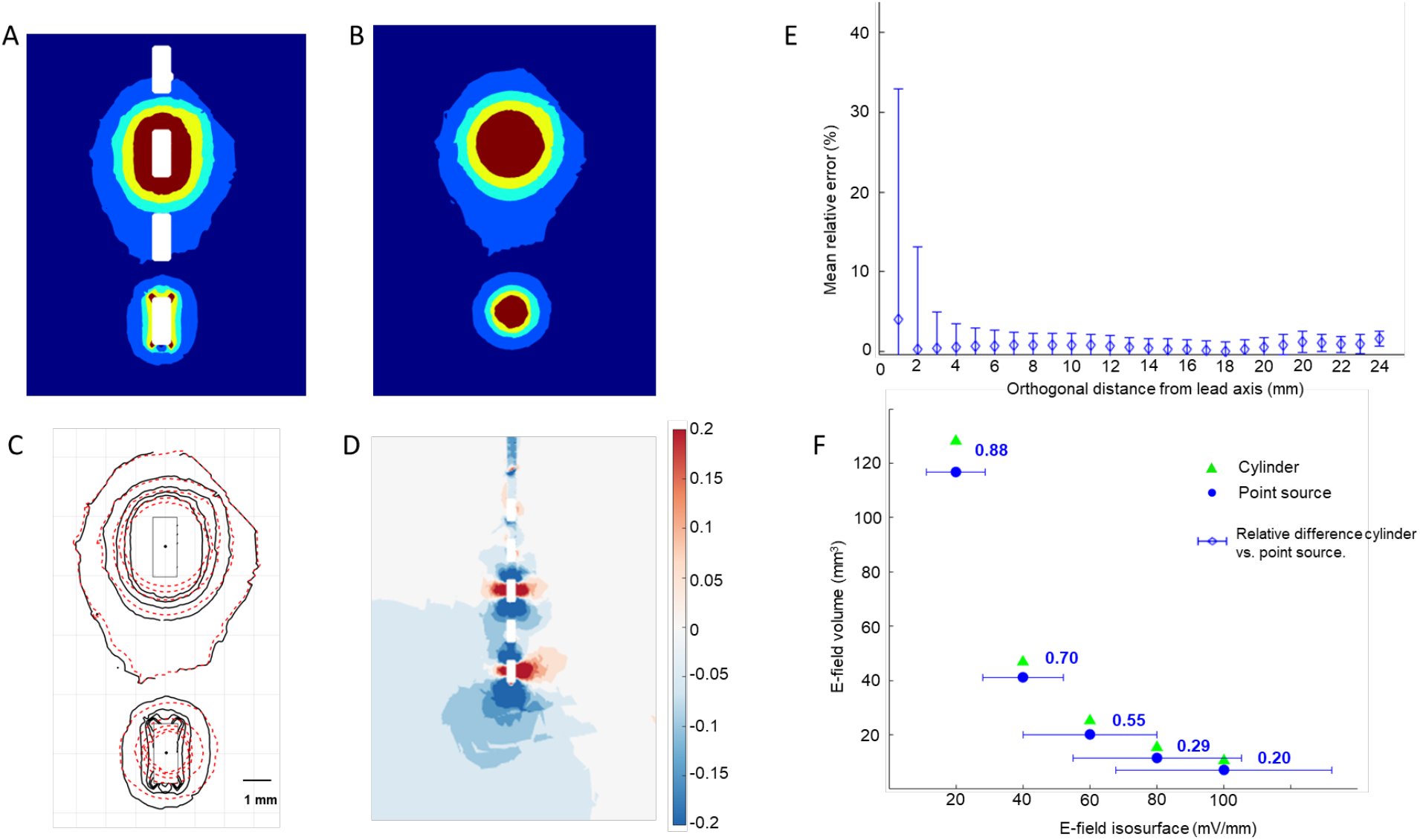
Realistic head model. A) Electric field distribution at the plane crossing the electrode for a non-adjacent configuration injecting ±1 mA with the cylindrical model of the electrode, and B) the point source. C) Isofield lines (20, 40, 100 mV/mm) for the realistic model (black lines) and point source approximation (red dashed lines). D) Relative error, calculated on the same plane, between the realistic and point source approximation. E) Mean relative error and standard deviation between the realistic model and the point source of the electric field as a function of the orthogonal distance from the electrode axis. F) Electric field volume within the 20, 40, 60, 80, 100 mV/mm isosurfaces for the three electrode models along the corresponding relative error bars and the Dice coefficient.

The isofield contours in Fig. 4c illustrate that the point source approximation overestimates the EField at distances closer than 3 mm from the electrode axis. The relative difference of the electric field Fig. 4d between the two electrode models also showed an overestimation of the point source approximation at the middle of active contacts, and an underestimation at the level of contact corners. In terms of relative error as a function of orthogonal distance, Fig. 4e showed the largest difference (~5%) at 1 mm from the electrode, and drastically decreased beyond 3 mm from the electrode axis. The EF volume difference ranged from 8 to 32 %, with lower values for low electric field magnitudes (Fig. 4f). The similarity between the two EF volumes was higher for lower EF magnitudes, with a Dice coefficient of 0.88 for 20 mV/mm and 0.2 for 100 mV/mm.

## Discussion

In this quantitative study, we have shown that both point and line source approximations can generate an electric field (EF) distribution comparable to the distribution obtained using a realistic cylindrical model of intracerebral electrode (SEEG) at distances of 3 mm or larger from the (cylindrical) electrode’s axis. In simplified head models, both approximations overestimated the EF, however the line source approximation achieved a better performance overall: not only it better matched the EF distribution shape obtained with the cylindrical model, but the Dice coefficient was also higher for all isofield values.

The graphs of the relative error at the plane crossing the electrode (panel E of Figures 1,2,3) support that none of the approximations could accurately reproduce the high current density at the edges between the active contact and the shaft insulating material [5-6]. Besides that, inactive contacts are not present in the line and point source approximations, the line source model considers a homogeneous current density along the entire line, while this magnitude actually varies along the electrode surface (especially at the electrode-insulating material interface) in the realistic model. Thus, although the field distribution is shaped more similarly to the realistic model, the assumption of a homogeneous current density prevents this approximation from correctly estimating the field magnitude. One possible direction for improvement would consist in setting non-homogenous current density functions along the source line approximating each contact in the line source approximation, and comparing the performance of such current density functions.

This difference between models was also observed in a study by Zhang and Grill [7]. Despite the fact that the authors used another variable (second spatial derivative of the potential) to test the point source approximation, the conductor-insulation interface was one of the sites where large differences were obtained between the models. In this regard, our results replicate and support further those reported in [7]. At the level of inactive contacts, we found small differences between the realistic model and its approximations, since we set the cylinder boundaries as insulating material.

In the case where the tilted electrode was crossing several layers with different conductivities, the point source model resulted in a slightly lower relative error for distances close to the electrode (Fig. 3g). Nonetheless, the Dice coefficient was still higher for the line source approximation, even for high isosurface values (> 100 V/m). Importantly, the field isosurfaces and corresponding volumes are commonly used to assess the spatial coverage of the field during brain stimulation [8]. Also, the volume comparison showed a larger relative difference between the point approximation and cylindrical model, especially for higher volumes (section H of Figures 1,2,3).

In this study, we assumed that SEEG electrodes were placed mostly in grey matter, and simulated this explicitly in the spherical model where active contacts were entirely surrounded by grey matter. The toy model, in contrast, intended to test the approximations in the presence of conductivity discontinuities, i.e., an inhomogeneous medium. Modeling the volume conductor as isotropic may correctly estimate the electric field as long as the contacts are located in grey matter, which has relatively uniform electrical properties [9]. Notably, it should be emphasized that the characteristic anisotropy of white matter may influence the field volume estimated by approximations. Notwithstanding these limitations, results indicate that the estimated field spatial distribution can provide essential information regarding the possible electric field impact on brain regions located in the vicinity of the stimulated region.

Finally, results using the point source showed that electrode geometry has an influence on E-field magnitude only at short distances (less than 3 mm from the electrode axis). We argue that this limitation is a not a significant one, since estimating the electric field too close to the electrode might be neither useful (since the volume of tissue close to the electrode is very small compared to the total volume impacted by the electrode contacts) or relevant (since a layer of gliosis surrounds the electrode, and develops over a few days following insertion of the electrode). Therefore, our results bring further support to the notion that approximating electrode geometries with simple sources (0D, 1D) can still provide useful quantitative characterizations of *in situ* Efields, while providing a considerable gain in terms of computation time, but also in terms of model implementation and use.

## Conclusion and perspectives

In this modeling study, we used a realistic model of SEEG electrodes as a ground truth to provide elements of comparison supporting the use of reliable approximations for fast computation of the electric field during neurostimulation performed using those electrodes. For distances close to the electrode, and as expected, the error of electrode model approximations increased drastically. Therefore, unless an especially accurate estimation of the E-field near the contacts is needed (up to 2 mm), the line source approximation is a suitable model for estimating the electric field generated during neurostimulation protocols in patient-specific models. We argue that the line source approximation can be used in confidence for most electric field dosimetry applications involving intracranial SEEG electrodes.

In terms of future prospects, data from the tested approximations could be used to validate their accuracy on clinical SEEG signals acquired during electrical stimulation. Typically, during pre-surgical evaluation of drug-refractory epilepsy, pulsed stimulation (intensity typically < 5 mA) is performed in clinical routine to identify epileptogenic regions. Knowing the stimulating contacts, the actual spatial profile of the E-field could be calculated during each stimulation pulse, and compared with the simulated E-Field both in terms of magnitude and distribution. It should be kept in mind that such endeavors might involve further developments in terms of biophysics: while we considered an ideal, homogeneous and isotropic medium in this study, which was purely resistive, we might require to take into account capacitive effects, that can be induced by the electrode-electrolyte interface for example [10]. Tissue properties that underlie the characteristics of the electrode-electrolyte interface include, but are not limited to, CSF infiltrations after SEEG electrode insertion, or gliosis tissue gradually encapsulating the electrode over the days following surgery. A dedicated biophysical study could focus on investigating the contributions of such factors on the electrode-electrolyte interface on the one hand, and on model accuracy when comparing with clinical data on the other hand.

## Methods

### SEEG electrode model

The realistic SEEG clinical electrode model consisted of an array of cylindrical contacts (platinum-iridium, conductivity chosen as 1000 S/m), which were 2 mm long and 0.8 mm in diameter, as shown in Fig. 5 separated by 1.5 mm of insulating parts (conductivity chosen as 0.001 S/m). In the case of the 0D (point source) approximation, sources were located at the center of mass of electrode contacts; while for the 1D (line source) approximation, source lines were placed along their central axis (as shown in blue in Fig. 5). A floating potential boundary condition was imposed at the surface of active contacts with a current of ± 1 mA. Inactive contacts were set as electric insulation, according to a recent study where it was shown that metallic materials act as electric insulators when exposed to electric fields up to 100 V/m [11]. The point and line approximations (0D and 1D, respectively) were set respectively to a point or line current source of ± 1 mA.

**Figure 5.**
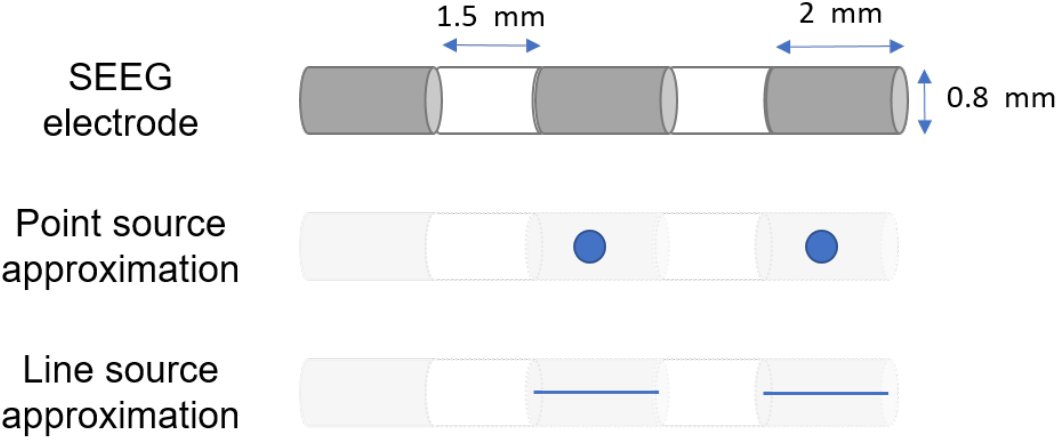
Schematic representation of the electrode geometries used to simulate the electric field induced by SEEG electrodes. In general, clinical SEEG electrodes include 10 to 15 contacts (gray color; length: 2 mm, diameter: 0.8 mm) separated by insulating material (white color; length: 1.5 mm). Here, the total number of contacts used were 14; the three model geometries were meshed in Comsol resulting in 2,327,559; 1,013,248; and 1,014,248 tetrahedral elements for the realistic, point source, and line source models, respectively.

### Volume conductor model - Spherical and toy model

Two types of head models were used to quantify the performance of our electrode model approximations: 1) a spherical model with 6 concentric spheres (Fig. 7a), and 2) a toy model representing tissue layers with concentric cubes with a simplified sulcus representation (Fig. 7b) based on [12]. For both models, the outer layer corresponded to the scalp with a conductivity of 0.33 S/m, followed by the skull set to 0.008 S/m, the cerebrospinal fluid (CSF) set to 1.79 S/m, and the grey and white matter with a conductivity of 0.4 and 0.15 S/m, respectively [12]. The spherical model had an extra inner layer for grey matter, considering that for some SEEG explorations deep electrode contacts can be placed in subcortical structures whose conductivity is similar to that of neocortex.

Domains’ geometries were built using the predefined elementary shapes in Comsol Multiphysics v5.6 (Comsol AB, Stockholm, Sweden). The electric field magnitude was computed by solving the Laplace equation using a steady-state approximation in Comsol: ∇·(σ∇V) = 0, where V is the electric potential (V) and σ is the electrical conductivity (S/m). The choice of the Laplace equation implies the following assumptions: 1) electrical stimulation is typically performed at low frequencies for pre-surgical exploration in drug-refractory epilepsy (typically < 100 Hz), justifying the quasi-static approximation which is considered as valid up to the kHz range; and 2) electrical conductivity was chosen as constant for a given tissue type, assuming a homogeneous and isotropic medium.

### Volume conductor model - Realistic model

To create a realistic model of an implanted SEEG electrode, we started by designing the 3D geometry of the lead in Comsol (version 5.3a). The positions of the centers of active contacts were obtained from segmentation (using the software Gardel [13]) of a head CT scan registered to the T1w-MRI that was used to create the head model. Each active contact was then represented as a cylinder with a radius of 0.4 mm and a length of 2.0 mm centered in each of those positions. These cylinders were then aligned with other cylinders (same radius), representing the SEEG shaft positions without active contacts. The SEEG electrode was prolonged from the last position, so that it finished 15 mm outside the scalp of the head model. The burr-hole created during the implantation was modeled as a larger cylinder with a diameter of 1.5 mm, surrounding the entire shaft. This geometry was exported from Comsol as a *stl* file. It was then added to the surface meshes of the head model (whose creation was described previously) using the library Pymesh (https://pymesh.readthedocs.io/en/latest/). Using Pymesh, a Boolean symmetrical difference was performed between the surface meshes of the tissues in the head model and those in the SEEG lead. The resulting surface was inspected for intersections and exported into Matlab (v2018b, www.mathworks.com), where a volume mesh was created using the TetGen library (https://people.math.sc.edu/Burkardt/examples/tetgen/tetgen.html) as implemented in Iso2mesh (http://iso2mesh.sourceforge.net/). The mesh comprised approximately 4 million tetrahedral finite elements with an average element quality of 0.57. The volume mesh was imported into Comsol, where appropriate electrical conductivities were assigned to each region: the head models were assigned to isotropic conductivity values as mentioned before, whereas the SEEG lead was assigned to a low conductivity, following [11]. Regarding the burr-hole in the skull and scalp, it was assigned to a conductivity of 1.79 S/m (same as CSF). The burr-hole in WM and GM was considered negligible and represented with the same conductivities as those tissues. The final head model with the added SEEG electrode is shown in Fig. 6.

**Figure 6:**
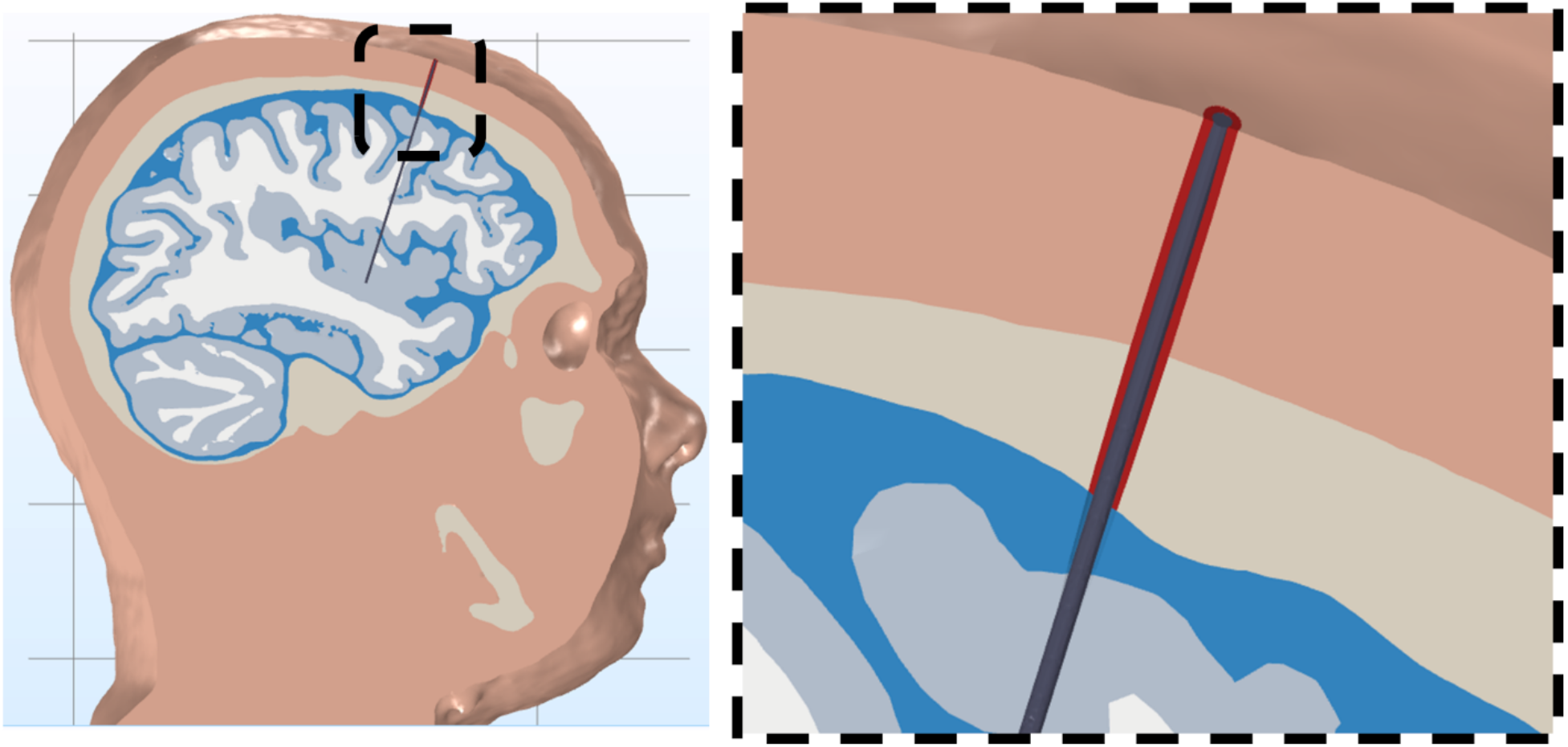
Sagittal view of the head model with the added SEEG lead. The caption shows, in more detail, the burr-hole that surrounds the SEEG lead in the scalp and skull (in red).

**Figure 7.**
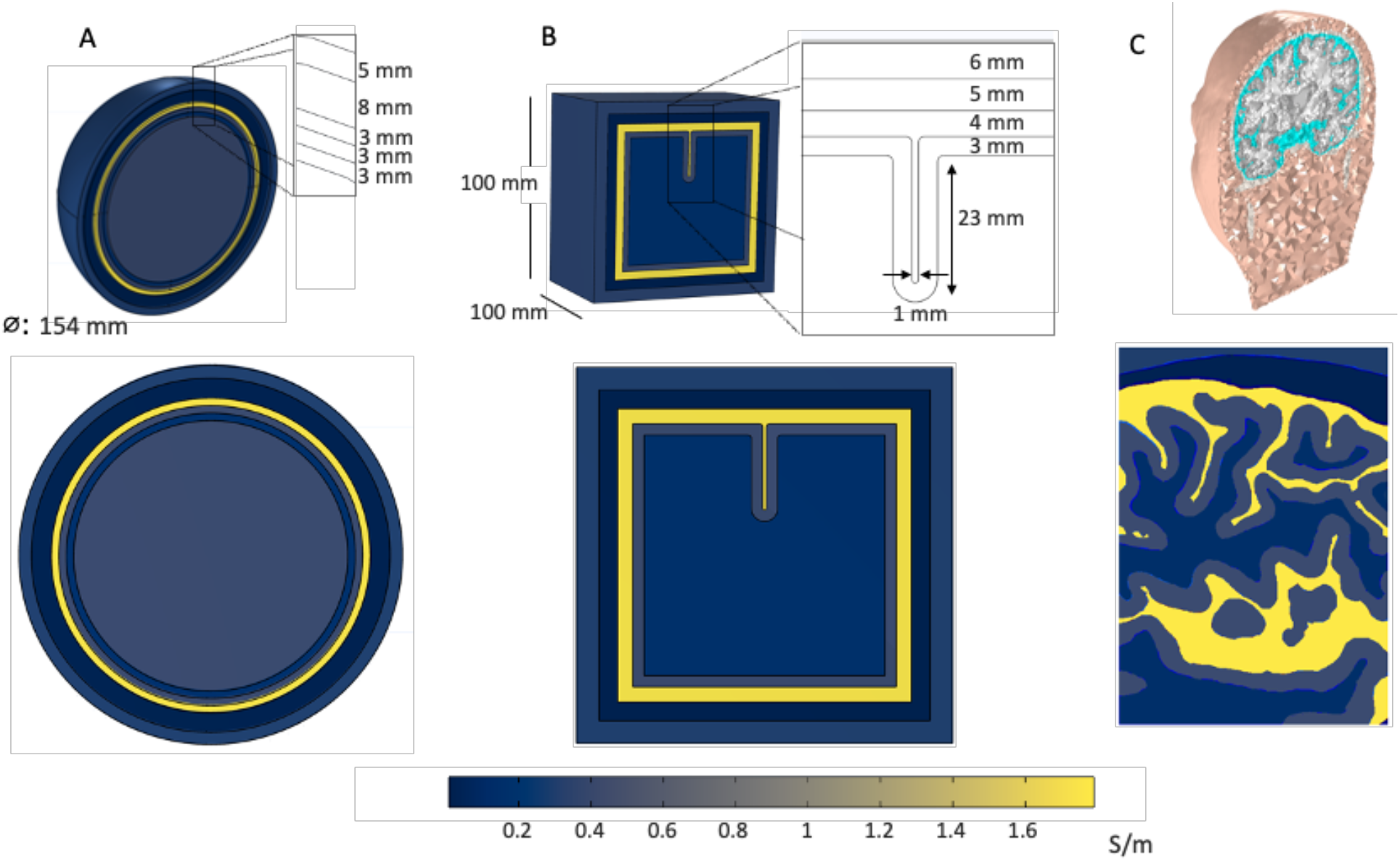
Volume conductor geometries used to test different approximations for SEEG electrode geometry. A. Spherical model where the grayish inner sphere and third layer correspond to gray matter with a conductivity of 0.4 S/m; second layer, in blue, represents the white matter (0.15 S/m); CSF is represented by the yellow layer (1.79 S/m), skull by the navy blue layer (0.01 S/m), and the scalp by the bluish outer layer (0.33 S/m). B. Toy model with a simplified representation of a sulcus, in the same color code, where the inner cuboid in blue, corresponds to the white matter; second layer, in grayish, represents the gray matter. C. Coronal view of the realistic head model (top) and sagittal cut (bottom) showing CSF, scalp, gray and white matter electrical conductivity using the same color code.

Electric field distribution was computed for two configurations: 1) adjacent electrode contacts, and 2) non-adjacent electrode contacts. In the case of the spherical head model, active contacts were those close to the tip of the shaft (Fig. 8a). For the toy model, the electrode was placed either parallel (Fig. 8b) to the sulcus, or tilted (Fig. 8c), crossing the sulcus and placing active contacts at the sulcus level to increase geometry complexity and challenge the robustness of the electrode model performance. In the realistic head model case, the electrode crossed several layers of tissue, as presented in Fig. 8d.

**Figure 8.**
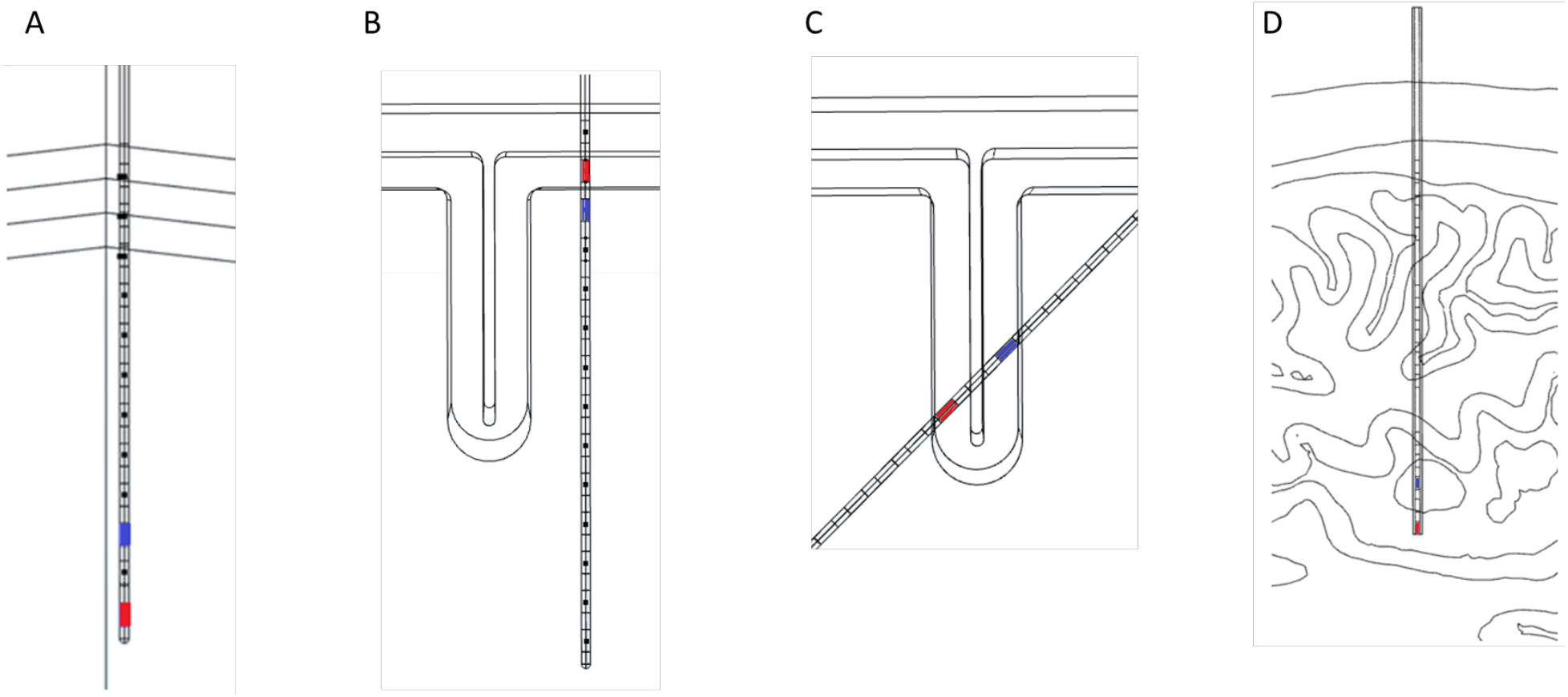
Positioning of active contacts. A) C1 and C3 contacts used as anode/cathode in a non-adjacent configuration placed in the inner layer of the spherical model, which corresponds to gray matter. B) Electrodes placed parallel to the sulcus in the toy model with adjacent active contacts with C12 placed in white matter and C13 in gray matter. C) Tilted trajectory where the electrodes cross the sulcus in a non-adjacent configuration placing the active contacts, C5 and C7, in grey matter. E) Realistic head model and non-adjacent configuration of active electrodes with C1 in the CSF and C2 in grey matter.

To evaluate the similarity of the electric field estimated by each model, we calculated the Dice coefficient for the binarized regions (<10, 20-40, 40-60, 60-80, and >100 mV/mm) of the electric field according to the equation:

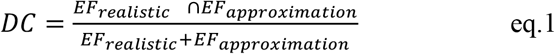

The average relative error was computed as a function of the distance from the electrode to quantify the performance of the two approximations for different distances, and estimate the regions in which those can be used reliably. A 3D grid with a resolution of 0.25 mm was used to extract the electric field magnitude computed in Comsol; for each point of the grid, we computed its distance to the electrode axis. These distances were then rounded to millimeters to pool grid points and calculate the mean error at each distance range. We also computed the volumes enclosed by the field isosurfaces for various field magnitudes. This was performed for all models using Comsol built-in functions, with the goal of providing an additional metric to compare the different models. Isocontours of 20, 40 and 100 mV/mm at the plane crossing the electrode were also visualized superimposing a 2-D grid with side lengths of 1 mm, centered on the electrode axis.

## Acknowledgements

The authors would like to acknowledge Denys Nikolayev for the line source approximation proposal.

